# DeltaNMF: A Two-Stage Neural NMF for Differential Gene Program Discovery

**DOI:** 10.64898/2026.01.22.701049

**Authors:** Anish Karpurapu, Charles Gersbach, Rohit Singh

## Abstract

Non-negative matrix factorization (NMF) is a foundational dimensionality-reduction method in single-cell transcriptomics, valued for its interpretable gene programs. However, in case-control settings common in perturbation and disease research, standard NMF conflates quantitative shifts in program usage with qualitative emergence of novel programs, obscuring whether observed differences reflect altered activity of shared programs or genuinely new cellular states. Here, we present DeltaNMF, a neural network-based reformulation of NMF that enables flexible biological priors and structured multi-stage fitting. We first demonstrate how our framework incorporates foundation model-derived gene similarities through graph-Laplacian regularization, improving program coherence by 2.4-fold on protein interaction networks. Building on this flexibility, we introduce a two-stage architecture that explicitly separates baseline programs (learned from control cells) from case-specific programs, disambiguating altered usage from novel program emergence. Our GPU-accelerated implementation achieves 20× speedup over consensus NMF while maintaining comparable accuracy. On synthetic data with known ground truth, DeltaNMF correctly identifies which programs are new versus shifted. Applied to coronary artery disease (CAD) Perturb-seq data, DeltaNMF isolates disease programs including the cerebral cavernous malformations pathway identified by Schnitzler et al., while clearly distinguishing them from baseline endothelial programs with altered activity. Our neural network formulation of NMF opens new directions for incorporating diverse biological priors into interpretable single-cell analysis, providing a principled framework for differential program discovery in case-control studies.

## 1 Introduction

A central challenge in understanding cellular heterogeneity from single-cell transcriptomics is extracting interpretable structure from high-dimensional single-cell RNA-seq (scRNA-seq) data. Dimensionality reduction methods like principal component analysis (PCA) and non-negative matrix factorization (NMF) are foundational tools for identifying patterns in these data. NMF is particularly attractive because the non-negativity of both factors (*W*, representing gene programs) and loadings (*H*, representing program activity per cell) offers biological interpretability. Gene programs derived from NMF factors have been found to correspond to biological processes, while loadings indicate each program’s activity in individual cells. This interpretability has driven widespread adoption [2, 20] and also adaptation: cNMF [12] is a standard tool for Perturb-seq analysis; LIGER and coupled NMF [5, 23] use NMF for multimodal integration; scInsight, which combines NMF programs across samples [18]. Here we focus on an increasingly common setting of NMF use: case-control studies, where researchers compare disease versus healthy tissue [6, 11], perturbed versus control cells [16, 19, 22], or drug responders versus non-responders [8, 14].

Current NMF approaches face two critical limitations: a) in case-control settings, a difficulty distinguishing program-level changes, and b) weak biological coherence of inferred programs. Across case-control cell groups, gene programs can differ *quantitatively*, where the same program exists in both populations but shows altered loading, or *qualitatively*, where an entirely new program emerges in the case population. The standard NMF approach fits a global set of factors on pooled data, then tests for differential loadings between groups, conflating these scenarios. A genuinely new case-specific program may be missed and its weights distributed as fractional loadings across multiple control programs, thus obscuring whether we observe a quantitative shift or qualitatively new cellular states. This ambiguity is exacerbated in settings where the case or control cells outnumber the other (e.g., common in Perturb-seq)—the global NMF fit naturally emphasizes patterns explaining the larger population. Subsampling can address this but risks losing statistical power.

The second limitation concerns biological coherence: locally-optimal NMF solutions often produce programs with redundant or unclear functional signatures, with multiple programs capturing overlapping pathway aspects rather than crisply delineating distinct processes. One cause is optimization complexity—finding the global optimum for *min*_*W*_, 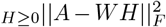 is NP-hard, so iterative algorithms converge to local optima. Another cause is the noise and sparsity of single-cell RNA-seq itself. cNMF’s strategy of multiple random restarts plus consensus clustering mitigates this but requires substantial computation.

We reformulate NMF as a neural network optimization that encodes biological structure directly into the fitting procedure, enabling both explicit case-control modeling and biologically-informed regularization (Fig. 1). One of our insights is to leverage ReLU-like activations to enforce non-negativity of *W* and *H* in an end-to-end differentiable formulation. This enables flexible loss functions, GPU acceleration, and principled incorporation of biological priors. Our first innovation regularizes gene program coherence using gene-gene similarity from scGPT and TranscriptFormer, single-cell foundation models [3, 17]. By encouraging genes within a program to agree with the foundation model’s learned semantic space, we produce factors with sharper functional delineation. Our second innovation introduces a two-stage case-control architecture: we learn a compact program dictionary on control cells alone, then fit additional case-specific programs for case cells while also keeping control programs active. This architecture explicitly separates control programs with changed loadings from entirely new case-specific programs.

**Fig. 1:**
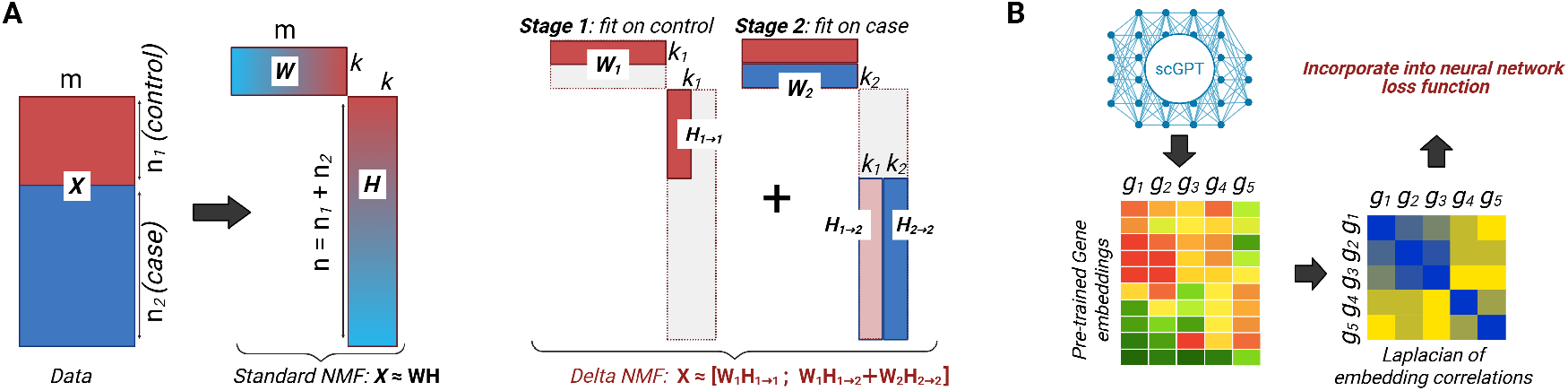
DeltaNMF overview and architecture. **(A)** Standard NMF fits a single set of programs to pooled case-control data, conflating quantitative changes (altered program usage) with qualitative changes (new programs). DeltaNMF uses a two-stage architecture: Stage 1 discovers baseline programs from control cells; Stage 2 fits additional case-specific programs while keeping baseline programs active, explicitly separating new programs from shifted usage patterns). **(B)** DeltaNMF uses embeddings from transformer models in its neural network loss function to refine gene loadings on gene programs.

We validate DeltaNMF on synthetic data where ground truth is known, demonstrate superior biological coherence through foundation model regularization, and apply it to uncover disease mechanisms in coronary artery disease (CAD) and Alzheimer’s disease (AD) Perturb-seq datasets. In both Perturb-seq screens, CRISPR interference is used to perturb genes nominated by disease GWAS in disease-relevant cell types (endothelial cells for CAD, primary human macrophages as a microglial proxy for AD), and single-cell RNA-seq is used to read out the resulting transcriptional programs. On synthetic benchmarks, DeltaNMF accurately recovers the ground-truth programs, matching cNMF’s accuracy at ∼20× its runtime and ∼200-400× greater computational efficiency. Foundation model regularization produces factors with over 2.4-fold improvement in STRING network consistency and a 2-fold increase in non-redundant GO terms compared to unregularized baselines. In Schnitzler et al.’s CAD study, the authors identified cerebral cavernous malformations (CCM) pathway involvement as a driver of atherosclerosis [19], a finding also identified by DeltaNMF. In the AD Perturb-seq dataset, DeltaNMF isolates a distinct lipid–lysosome program that links APOE-mediated lipid uptake with coordinated activation of lysosomal cathepsins, mirroring the dysfunctional plaque-processing phenotype observed in human microglia.

## 2 Methods

### 2.1 Problem Formulation

Non-negative matrix factorization (NMF) decomposes a non-negative data matrix 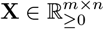 (with *m* genes and *n* cells) into the product of two non-negative matrices **X** ≈ **WH** where 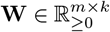 contains *k* gene programs and 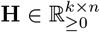 contains the usage of each program in each cell. The standard NMF objective minimizes the Frobenius norm of the reconstruction error:

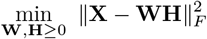

#### Consensus NMF (cNMF)

The NMF optimization problem is non-convex and NP-hard in general. Individual runs can be unstable: different random initializations of **W, H** can converge to diverse local minima composed of distinct programs. Consensus NMF (cNMF) addresses these issues by averaging the gene program structure over many independent factorizations [12]. In practice, one performs *r* runs with different seeds, pools the *r* × *k* program columns, clusters them by similarity, and aggregates each cluster to form a consensus **W**. The corresponding **H** is then refit by nonnegative least squares against the fixed consensus dictionary. This view can reduce variance, mitigate spurious programs, and reflects reproducible structure. While it does fall short of providing principled guarantees, cNMF is now a popular NMF implementation and often used in Perturb-seq gene program analyses [12].

### 2.2 Overview of DeltaNMF

Our starting observation is that NMF can be reformulated as a GPU-friendly optimization problem. We parameterize **W** and **H** as learnable parameters in a single-layer neural network and enforce non-negativity through activation functions rather than explicit constraints. Specifically, with an elementwise ReLU activation *ϕ*(*x*) = max(0, *x*), the reconstruction is:

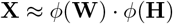

We supervise with the mean-squared-error (MSE) loss: 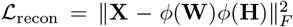 . After training, we report **W**_**eff**_ = *ϕ*(**W**) and **H**_**eff**_ = *ϕ*(**H**) as the learned factorization.

We use the Adam optimizer with a constant learning rate schedule [10]. Convergence is declared when the relative improvement in ℒ_recon_ falls below 10^−8^ or the epoch limit is reached. In practice, we found more stable convergence and comparable reconstruction losses using SoftPlus rather than ReLU (Supp. Fig. B.1). This is in line with general observations in neural network training that smoothening the ReLU non-linearity helps with training. We report the final matrices, **W**_**eff**_ and **H**_**eff**_ .

#### Batching and Initialization

The gene program matrix **W** ∈ ℝ^*m*×*k*^ typically fits in GPU memory even with limited VRAM (with *m* ∼ 20,000 genes and *k* ∼ 10–300 programs). However, **H** ∈ ℝ^*k*×*n*^ and **X** ∈ ℝ^*m*×*n*^ can be very large when *n* > 100,000. We therefore process **X** and **H** in batches of size *B*. For batch *i*:

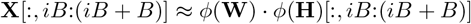

The batch size *B* can be calibrated to available GPU memory. An epoch of DeltaNMF corresponds to a single pass over all observations of **X**. Like standard NMF solvers, our GPU-based formulation also finds local optima. Following cNMF, we do multiple short runs and our final reported parameters are from a DeltaNMF run initialized with consensus parameters (Section A.1).

### 2.3 Customizing the loss function: Foundation Model-Informed Regularization

A key advantage of our neural network formulation is that multiple loss terms can now be added to the objective function and optimized using standard neural network training machinery. This was previously difficult and cumbersome. For instance, incorporating additional constraints into Yang and Michailidis’ integrative NMF [24] objective required substantial algorithmic development. To demonstrate this capability, here we introduce an additional loss term to encode a biological prior.

Specifically, we seek to regularize NMF factors in settings where limited data might lead to noisy program estimates. Our intuition is that, aggregated across millions of transcriptomes, there exists a baseline pattern of gene co-activity that programs in any sample should “shrink” towards. To regularize toward this baseline, we use gene embeddings from scGPT, a single-cell foundation model trained on 33 million cells across diverse tissues and conditions [3].

The overall objective combines reconstruction error with foundation model regularization:

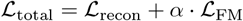

where ℒ_FM_ is the added foundation model-informed regularization loss and *α* ≥ 0 is the regularization weight. We compute ℒ_FM_ by obtaining pre-trained gene embeddings from scGPT, computing gene–gene similarities and encoding that into a program-consistency objective.

#### Gene Similarity Graph

Let **E**_fm_ ∈ ℝ^*m*×*d*^ be pre-trained gene embeddings from a single-cell foundation model (for scGPT, *d* = 512) [3]. We build a symmetric similarity matrix **S**_*E*_ with entries

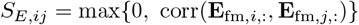

where corr(·, ·) denotes Pearson correlation. Let **D**_*E*_ be the diagonal degree matrix with **D**_*E,ii*_ = ∑_*j*_ **S**_*E,ij*_.

#### Graph Laplacian Regularization

We use the graph Laplacian **L**_*S*_ = **D**_*E*_ **S**_*E*_ to penalize roughness of program loadings across the gene similarity graph:

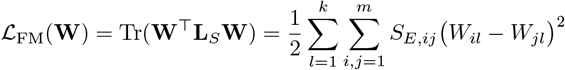

This term promotes programs where genes that co-load are functionally related according to the foundation model’s learned semantic space. When genes *i* and *j* have high similarity *S*_*E,ij*_, the penalty encourages their loadings *W*_*il*_ and *W*_*jl*_ to be similar within each program *l*.

#### Interpretable hyperparameter specification

The scale of loss terms depends on the dataset, making it difficult to specify *α* consistently across different analyses. To enable an interpretable and stable choice, we first compute the relative scale of the reconstruction and regularization losses, then express *α* as a multiplier on a user-specified baseline strength *α*_0_:

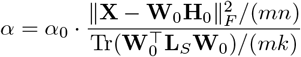

where (**W**_0_, **H**_0_) come from the consensus initializer. This adaptive scaling makes *α*_0_ interpretable and invariant to global data scaling and comparable across datasets.

### 2.4 Two-Stage Case-Control Architecture

For case-control studies (e.g. disease vs. healthy, perturbed vs. control), standard NMF can conflate quantitative shifts in program usage with qualitative emergence of new programs. We introduce a structured two-stage approach where we separate these effects by first discovering baseline programs on control cells, then fit additional case-specific programs while keeping the baseline active.

Let 𝒢 be the set of genes and 𝒞 = 𝒞 ^(1)^ ∪ 𝒞 ^(2)^ the set of cells, partitioned into controls 𝒞^(1)^ and cases 𝒞^(2)^. We form count matrices **X**^(1)^ (control) and **X**^(2)^ (case), with *m* = |G|, *n*_1_ = |𝒞^(1)^|, *n*_2_ = |𝒞^(2)^|:

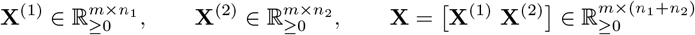

#### Stage 1: Baseline program discovery (controls)

Let 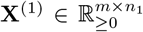 denote the control matrix. We learn *k*_1_ baseline programs by

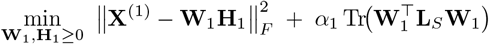

where 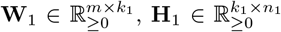, and **L**_*S*_ is the graph Laplacian built from scGPT gene similarity. We use the initialization strategy described earlier, then refine with the FM loss regularization, weighted by *α*_1_. Denote the result by **W**_1_.

#### Stage 2: Augmented program discovery (cases)

Given the learned **W**_1_ from Stage 1 and the case matrix **X**^(2)^, we learn *k*_2_ additional programs 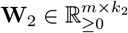 and case usages.

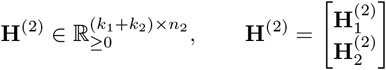

such that **X**^(2)^ ≈ [**W**_1_ **W**_2_] **H**^(2)^. We optimize the combined objective:

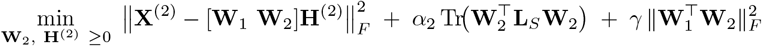

where *α*_2_ weighs the FM regularization in Stage 2. The orthogonality penalty 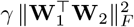 encourages case-specific programs to be distinct from baseline programs. To make this invariant to program scale, we implement this as the squared Frobenius norm of the cosine similarity matrix:

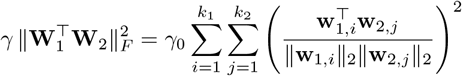

where **w**_1,*i*_ and **w**_2,*j*_ denote columns of **W**_1_ and **W**_2_ respectively. This formulation penalizes similarity between baseline and case-specific programs while remaining invariant to the magnitude of program loadings.

During Stage 2 we keep **W**_1_ fixed, updating **W**_2_ and **H**^(2)^ with column mini-batches of **X**^(2)^. Nonnegativity is enforced by elementwise softplus activation (*β* = 5.0) on the trainable parameters for **W**_2_ and **H**^(2)^. In practice, we set *α*_2_ = 0 if we expect the discovery of novel programs that the foundation model may not capture, for example, in rare disease states or context-specific perturbations. Stage 2 uses a residual-based initialization that first projects cases onto the control dictionary, factorizes remaining variance into candidate case programs, and refines them jointly (see A.2). The final **H**^(1)^ is re-fitted by fixed-factor NMF on the combined program matrix output from Stage 2: [**W**_1_ **W**_2_].

### 2.5 Implementation Details

Model training was primarily performed on NVIDIA A100-SXM4 (80 GB memory) GPUs. A dataset of 215k cells and ∼20k genes takes ∼2.5 hours for a full run of two-stage DeltaNMF. By comparison, standard cNMF required > 48 hours despite high-memory provisioning (10 cores, 150 GB RAM each). Consequently, DeltaNMF offers a ∼20× improvement in runtime and ∼200-400× speedup in resource-adjusted throughput.

## 3 Results

### 3.1 Validating the base DeltaNMF formulation

#### DeltaNMF accurately recovers synthetic data

To establish that our neural reformulation preserves NMF’s core capabilities, we first tested DeltaNMF on simulated data with known ground truth. Following Kotliar et al.’s Splatter-based simulation framework, we generated 15,000 cells comprising 13 distinct cell types (identity programs) and one activity program expressed across multiple cell types. The simulation included realistic technical noise including doublets and expression variability parameters from published organoid data [12, 25]. Applying vanilla DeltaNMF (Section 2.2; no additional regularization) to the pooled data, we recovered both the gene programs and their cell-specific activity patterns (Fig. 2A). All 14 programs achieved strong correspondence with ground truth (median Pearson *ρ* = 0.946, minimum *ρ* > 0.9), confirming that our neural formulation captures the same underlying structure as traditional NMF.

**Fig. 2:**
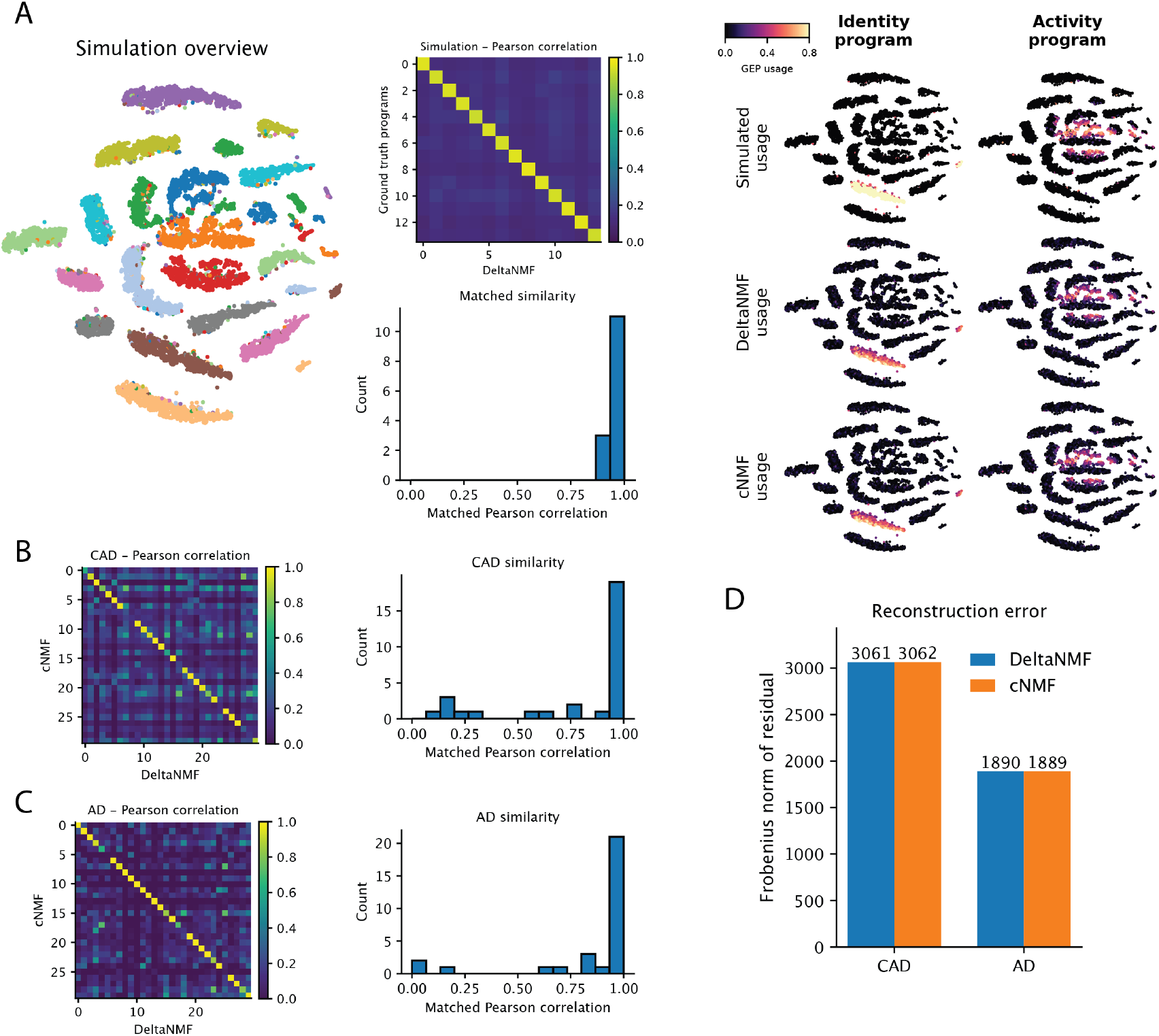
Validation of the base DeltaNMF formulation. **(A)** Synthetic data: t-SNE of 15,000 simulated cells, program-matching using Pearson correlation after Hungarian alignment, and usage maps for ground truth, DeltaNMF, and cNMF programs. **(B)** Coronary artery disease (CAD) scRNA-seq: correlation matrix between cNMF and DeltaNMF programs (k=30). **(C)** Alzheimer’s disease (AD) scRNA-seq: same comparison. **(D)** Reconstruction error (Frobenius residual) for cNMF and DeltaNMF across both datasets.

#### DeltaNMF accurately recovers programs in real-world data

We next verified that DeltaNMF produces comparable results to consensus NMF (cNMF) on real biological data. We used two Perturb-seq datasets: a coronary artery disease (CAD) study and an Alzheimer’s disease (AD) study (see Supp. Methods A.4). In this analysis, we limited ourselves to the baseline cells (i.e., with non-targeting guides) since we sought only the single-stage NMF fit. The CAD dataset comprised 214,449 immortalized endothelial cells, of which 5,506 received non-targeting guides. The AD dataset included 28,446 primary human macrophages cells; 2,184 cells received non-targeting guides. We ran cNMF and DeltaNMF (single-stage formulation) with k = 30 programs on the same 2,000 highly variable genes selected by cNMF’s overdispersion metric.

Both methods produced highly concordant programs. The program-by-program correlation matrices showed strong diagonals, indicating clear one-to-one correspondence (Fig. 2B-C). In CAD data, 20 of 30 program pairs achieved *ρ* > 0.8 (median *ρ* = 0.964); in AD data, 25 pairs exceeded this threshold (median *ρ* = 0.995). Reconstruction errors were nearly identical between methods (Fig. 2D), demonstrating that our neural optimization achieves the same explanatory power as established approaches while enabling the flexibility for subsequent innovations.

### 3.2 Foundation model regularization improves program coherence in DeltaNMF

Having established that our neural formulation recovers comparable programs to standard NMF, we next explored how it enables novel biological priors through flexible loss functions. We hypothesized that incorporating gene-gene similarity information from single-cell foundation models could improve the biological coherence of learned programs. To test this, we augmented the DeltaNMF programs learned above from CAD and AD control cells with a graph-Laplacian regularizer derived from scGPT embeddings. We activated this regularization only during the final 200 epochs to ensure it only acts as a refinement.

To build intuition for how regularization reshapes programs, we examined a representative case from the CAD dataset (Fig. 3A). Programs learned without (*α* = 0) and with (*α* = 1.0) FM regularization maintain the same global architecture—evidenced by the strong diagonal in their correlation matrix—but the latter induces specific, biologically meaningful shifts in gene loadings. The graph Laplacian acts as a smoothing operator that pulls functionally related genes together: genes with weak, inconsistent loadings in the baseline model converge to coherent high loadings in the regularized version. This results in programs that are less defined by isolated markers and more reflective of complete biological complexes.

**Fig. 3:**
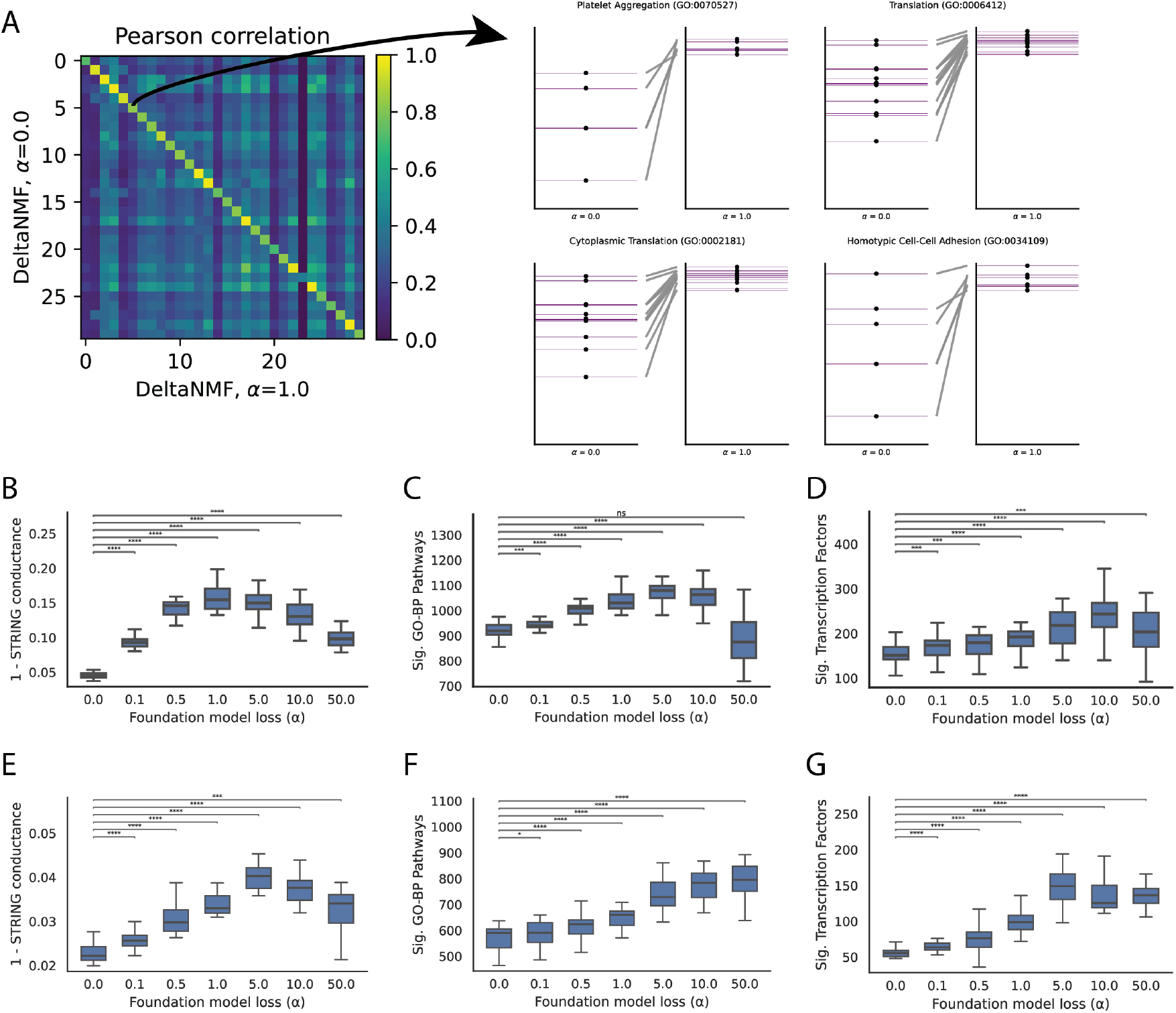
Foundation model regularization improves biological coherence of gene programs. Programs from CAD (B–D) and AD (E–G) control cells were refined using scGPT-based graph Laplacian regularization during the final 200 training epochs. Programs were evaluated across a range of regularization strengths (*α* ∈ [0, 50]) using bootstrap subsampling (80% of cells per replicate), and enrichment analyses used the top 300 genes per program. **(A)** Stability of program structure: correlation between baseline (*α* = 0) and regularized (*α* = 1) programs, and examples of gene-loading shifts. **(B–D)** CAD: STRING network cohesion (1-conductance), GO Biological Process pathway enrichment, and TRRUST transcription factor enrichment. **(E–G)** AD: the same three coherence metrics. Error bars show mean ± s.d. across 20 bootstrap resamples. Significance stars: ns for *p* ≥ 0.05, ^*^ for *p* < 0.05, ^**^ for *p* < 0.01, ^***^ for *p* < 0.001, ^****^ for *p* < 10^−4^.

We systematically evaluated this improvement across multiple dimensions of biological coherence. Testing regularization strengths from *α* = 0 (none) to *α* = 50 (strong) with 20 bootstrap resamples per condition, we found *α* = 1.0 optimally balanced coherence gains with reconstruction fidelity and recommended it as the default. At this setting, CAD programs showed significant improvements in functional cohesion (Fig. 3B-D). Protein-protein interaction connectivity, measured as network conductance within the STRING database, improved 243.5% (baseline: 0.046 ± 0.006 vs. regularized: 0.157 ± 0.019, *p* < 0.001). This three-fold increase indicates FM-regularized programs select genes substantially more tightly interconnected in protein-protein interaction networks, consistent with functional modules rather than arbitrary gene sets [21].

Beyond physical interactions, regularized programs better captured diverse aspects of biological organization. Gene Ontology (biological process) enrichment increased both in breadth (13.2% more significant pathways) and specificity (95.4% more non-redundant terms per program after semantic clustering, *p* < 0.001) [1]. Programs also showed enhanced transcriptional coherence (Supp. Fig. B.2), with 22.8% more enrichment for transcription factor target sets from TRRUST (*p* < 0.01), suggesting they better reflect coordinated regulatory modules [9, 13]. WikiPathways enrichment similarly improved by 20.6% (*p* < 0.001) [15]. Most strikingly, trait and disease relevance measured through GWAS gene enrichment from Open Targets L2G increased 180.9% (*p* < 0.001), demonstrating that improved functional coherence translates directly to stronger trait and disease associations [7]. These improvements remained consistent whether regularization was applied for 200 or 1000 epochs (Supp. Fig. B.4).

The AD dataset showed comparable improvements across all metrics: STRING cohesion improved 52.0%, GO-BP pathways 13.4% (Fig. 3E-G), as well as transcription factor enrichment 73.5% (Supp. Fig. B.2), WikiPathways 17.0%, and trait associations increased 267.2% (all *p* < 0.001). This consistency across independent datasets with different cell types (endothelial vs. macrophage) and disease contexts demonstrates that foundation model regularization provides a general strategy for enhancing program quality.

#### Robustness of regularization benefits

Several ablations confirm the stability of our approach. First, improvements were robust to regularization strength: while we recommend *α* = 1.0 based on optimal performance across most metrics, benefits were consistent for *α* values from 0.5 to 5.0 (Fig. 3B–G). Importantly, reconstruction error increased only marginally even at strong regularization (Supp. Fig. B.5), indicating that biological coherence gains do not sacrifice data fidelity. Second, the choice of foundation model proved flexible: TranscriptFormer embeddings yielded comparable improvements to scGPT (Supp. Fig. B.3), suggesting that benefits derive from general transcriptomic co-activity patterns learned by large-scale models rather than model-specific artifacts [17]. Together, these results establish DeltaNMF’s foundation model regularization as a robust, generalizable enhancement to NMF-based program discovery.

### 3.3 Two Stage DeltaNMF separates case-specific programs from baseline structure

#### Synthetic validation of program separation

The core innovation of DeltaNMF is distinguishing altered program usage from genuinely new programs. To test this directly, we designed a paired-program simulation where the ground truth explicitly separates these scenarios (Supp. Methods A.6). We generated 10 cell groups, each with a baseline program active in all cells and a case-specific program active only in case cells (Fig. 4A). Critically, baseline and case programs used completely disjoint gene sets, ensuring that any correlation between recovered baseline and case factors indicates unwanted signal bleeding rather than true biological overlap.

**Fig. 4:**
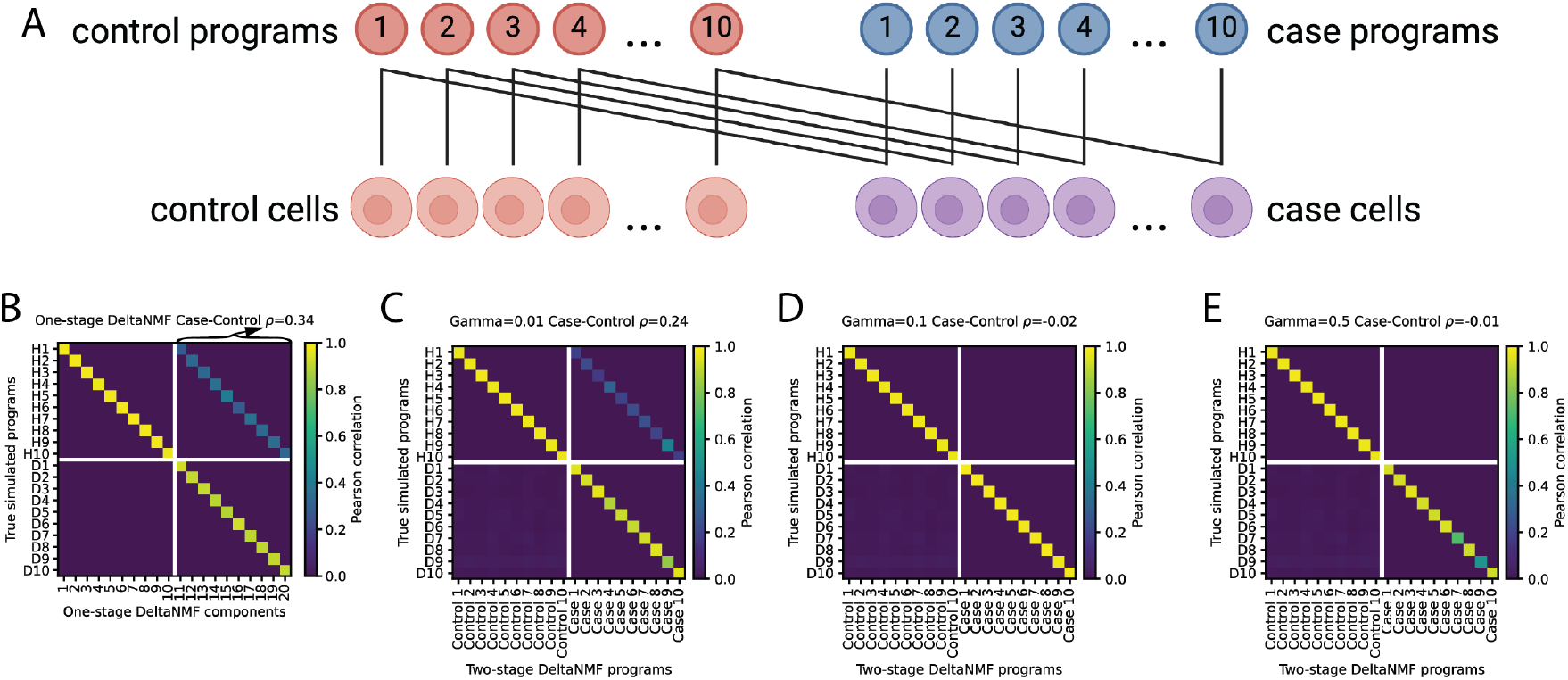
Two-stage DeltaNMF separates control and case programs in paired-program simulations. **(A)** Schematic of the simulation design. **(B)** One-stage baseline: correlation between inferred components and true programs after Hungarian matching. Control programs labeled H1-H10, case programs labeled D1-D10. **(C–E)** Two-stage DeltaNMF with increasing orthogonality penalties (*γ* = 0.5, 0.1, 0.01), showing reduced mixing between control and case programs.

We evaluated program recovery using correlation matrices between true and inferred programs after Hungarian matching. These matrices have four interpretable quadrants: the diagonal blocks show within-category recovery (control-control and case-case), while the off-diagonal blocks reveal cross-contamination. The upper-right quadrant specifically measures whether case-specific programs incorrectly capture control program patterns. We quantify this as *ρ*, the mean correlation between each true control program and its best-matching case program; values near zero are better.

Standard one-stage factorization on pooled cells recovers programs but conflates their origins (Fig. 4B). While the strong diagonal confirms program identification, the upper-right quadrant shows substantial bleeding (*ρ* = 0.34): case programs partially recapitulate control patterns even though they should be distinct.

The two-stage architecture with orthogonality penalty progressively eliminates this contamination (Fig. 4C– E). As we increase the orthogonality strength from *γ* = 0.01 to *γ* = 0.5, the upper-right quadrant clears (*ρ* decreases from 0.24 to -0.01), while program recovery along the diagonal remains intact. With *γ* = 0.5 or *γ* = 0.1, the upper-right quadrant became nearly empty, consistent with no systematic reuse of control programs by the case components. Importantly, in all settings the true case programs were still recovered along the main diagonal. Together, these simulations show the two-stage DeltaNMF architecture, with a moderate orthogonality term, cleanly separates new case-specific programs from baseline structure while preserving accurate recovery of both control and case programs.

#### CAD Perturb-seq validates separation of novel disease programs

Having demonstrated program separation in controlled simulations, we applied two-stage DeltaNMF to coronary artery disease (CAD) Perturb-seq data to test whether disease-associated transcriptional changes partition into altered baseline programs versus genuinely novel states. We configured the model with *k*_control_ = 30 baseline programs and *k*_case_ = 60 case-specific programs, selecting orthogonality penalty *γ* = 10^−3^ to balance program separation with reconstruction accuracy (Fig. 5A).

**Fig. 5:**
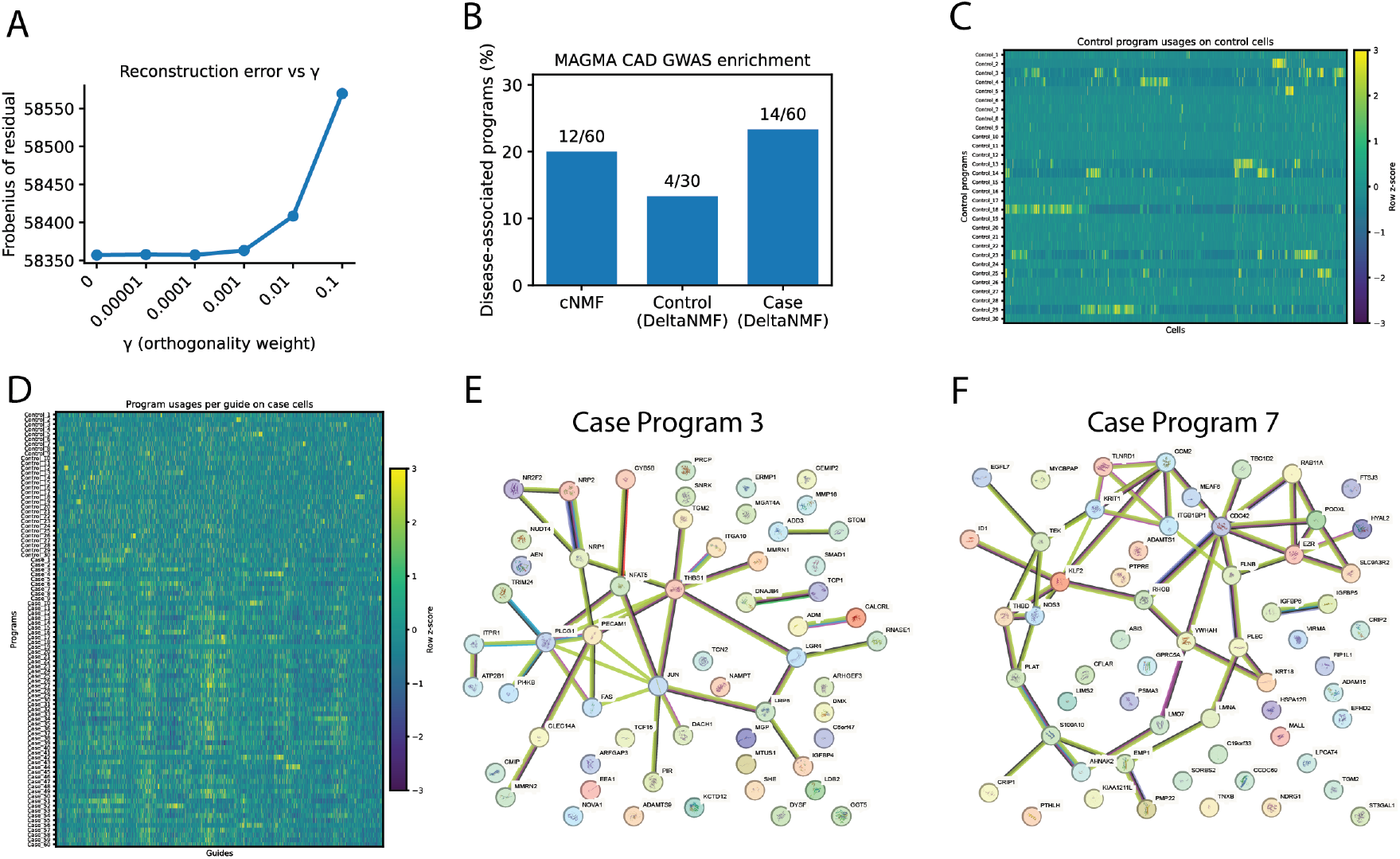
**(A)** Reconstruction error across orthogonality *γ* used to select *γ* = 10^−3^. **(B)** MAGMA-based CAD enrichment across cNMF programs and DeltaNMF control and case programs. **(C)** Control-cell usage of control programs. **(D)** Case-cell usage of control and case programs (collapsed by guide). For both heatmaps (C and D), usage values were row-standardized (Z-scored per program across cells), clipped to the range [−3, 3], and cells were hierarchically clustered by correlation distance. **(E-F)** STRING networks for two CAD MAGMA-associated case programs illus-trating angiogenic and CCM modules.

To identify which programs carry CAD genetic risk, we performed MAGMA enrichment analysis, which aggregates GWAS signal from SNPs to genes to programs while accounting for linkage disequilibrium and gene length [4]. This polygenic approach is ideal for testing our hypothesis: if CAD risk stems from quantitative shifts in normal endothelial function, signal should concentrate in baseline programs; if it involves qualitatively new cellular states, case-specific programs should dominate.

The results support DeltaNMF’s two-stage model with discovery of qualitatively new programs (Fig. 5B). While standard cNMF identified 12 CAD-associated programs (20% of 60 total), DeltaNMF revealed a notable asymmetry: only 4 of 30 baseline programs (13%) showed CAD enrichment (quantitative shift), whereas 14 of 60 case-specific programs (23%) carried significant signal (qualitative shift). This redistribution aligns with our conceptual framework. The few enriched control programs likely represent normal endothelial functions with altered activity in disease, while the numerous case-specific programs capture emergent pathological states. Usage patterns reinforce this interpretation: disease-relevant control programs show elevated expression broadly across perturbations, while case programs exhibit sharp, perturbation-specific activation (Fig. 5C-D and Supp. Fig. B.6).

Three case programs illustrate the distinct disease mechanisms captured by DeltaNMF. To identify each program’s upstream drivers, we linked programs to genetic perturbations by computing average program activity for each perturbed gene (Supp. Methods A.7).

Case Program 3 defines a pro-angiogenic state enriched for extracellular matrix remodeling (Fig. 5E). Central to this program are vascular stability regulators *CALCRL* and adrenomedullin (*ADM* ), guidance receptors *NRP1* and *NRP2*, and the matrix modulator *THBS1*. The top genetic drivers (*JUN, PLCG1*, and *FAS* ) delineate a signaling module associated with neovascularization, a critical process in plaque destabilization and intra-plaque hemorrhage.

Importantly, Case Program 7 recovers the cerebral cavernous malformation (CCM) pathway, independently validating the original study’s central finding (Fig. 5F) [19]. The program combines shear-stress responsive factors including master regulator *KLF2* and nitric oxide synthase *NOS3* with core CCM complex components *KRIT1* and *CCM2*. Notably, *TLNRD1* appears as both target and regulator, capturing the feedback loop essential for maintaining endothelial barrier function under hemodynamic stress.

Case Program 9 represents an inflammatory, atheroprone phenotype driven by NF*κ*B signaling (Supp. Fig. B.7). The program features extensive chemokine signatures (*CXCL1, CXCL8* /IL-8, *CCL2* /MCP-1) and inflammatory cytokines (*IL1A, IL1B, IL6* ) that recruit leukocytes, coupled with adhesion molecules like *ICAM1* and NF*κ*B feedback regulators (*NFKBIA, TNFAIP3* )—defining an activated endothelial state characteristic of atherosclerotic lesions.

By partitioning CAD risk into interpretable baseline and emergent programs, two-stage DeltaNMF reveals that disease pathology involves both quantitative dysregulation of normal endothelial functions and qualitative emergence of novel transcriptional states.

#### AD Perturb-seq confirms recovery of disease-relevant macrophage states

To further validate two-stage DeltaNMF’s ability to identify disease-specific programs, we analyzed the AD Perturb-seq dataset and compared recovered programs to those reported by Wang et al. [22]. Three case programs exemplify the pathological macrophage states captured by our approach (Fig. B.8).

Case Program 43 recapitulates the lipid-lysosome dysfunction central to AD pathology (MP1 in Wang et al.). This program links *APOE* —the strongest sporadic AD genetic risk factor—with lysosomal degradation machinery including acid lipase *LIPA* and cathepsins that process amyloid-beta aggregates. The co-expression pattern captures stressed lysosomes overwhelmed by lipid-protein debris from plaque phagocytosis, with genetic regulators *CD33* and *HEXB* directly modulating this degradative capacity.

Case Programs 48 and 7 capture complementary inflammatory states. Program 48 represents hyper-metabolic activation (MP5), dominated by ribosomal proteins and chaperones required for rapid cytokine production. Program 7 identifies an interferon response signature (MP2) featuring canonical ISG genes, suggesting sterile inflammation triggered by damage-associated molecular patterns from neurodegeneration. The independent recovery of these established AD-associated programs, particularly the *APOE* -linked lipid-lysosome axis, demonstrates that two-stage DeltaNMF reliably identifies disease-relevant transcriptional states across different biological contexts and cell types.

## 4 Discussion

Case-control studies are fundamental to single-cell biology, yet standard NMF approaches face a critical interpretive challenge: when programs differ between conditions, we cannot distinguish whether baseline programs are simply used more (quantitative change) or entirely new programs have emerged (qualitative change). DeltaNMF addresses this challenge through three synergistic innovations. First, our two-stage architecture explicitly separates baseline programs from case-specific ones, resolving the conflation. Our simulations demonstrate that standard approaches inherently blend case signals into control programs even when ground truth programs are completely distinct, while two-stage fitting with orthogonality constraints achieves near-perfect separation. This disambiguation is crucial for Perturb-seq and disease studies where distinguishing dysregulated baseline processes from emergent pathological states guides mechanistic understanding. This is exemplified by our recovery of the CCM pathway as a distinct case-specific program in CAD, validating the original study’s findings while clarifying its disease-specific nature.

We demonstrate the first—to our knowledge—successful integration of single-cell foundation models into interpretable dimensionality reduction methods like NMF. By incorporating gene-gene similarities from scGPT through graph-Laplacian regularization, we achieve strong improvement in protein-protein interaction coherence and disease gene enrichment. The consistency across different foundation models (scGPT and TranscriptFormer) suggests we’re capturing fundamental co-expression patterns learned from millions of cells, patterns that will only improve as foundation models advance.

Both these innovations rely on our neural network reformulation of NMF. Where previous methodological extensions required substantial algorithmic development, we can now incorporate arbitrary differentiable objectives through standard backpropagation. This democratization of method development, combined with 20× speedups from GPU acceleration, makes sophisticated analyses accessible to the broader community. Importantly, each innovation reinforces the others: the neural framework enables both the two-stage architecture and foundation model integration, which would be cumbersome to implement with traditional coordinate descent solvers.

DeltaNMF has some limitations. The two-stage architecture assumes discrete case-control separation, making it less suitable for continuous processes like differentiation trajectories. We assume control cells represent “pure” baseline states, which may not hold if controls harbor subtle perturbations. The orthogonality penalty, while effective for separation, might overly restrict programs that legitimately share components. Finally, while GPU requirements are increasingly standard, they may limit accessibility for some users.

Looking forward, the neural framework opens exciting extensions. Semi-supervised formulations could incorporate perturbation metadata directly into the loss function, ensuring programs map to specific genetic drivers. Multi-condition studies could use hierarchical architectures to model shared and condition-specific programs across many states. DeltaNMF is another in the line of approaches that reimagine classical methods through modern deep learning infrastructure to resolve long-standing analytical challenges while maintaining the interpretability essential for biological insight.

## Code and Data availability

The DeltaNMF code is publicly available at https://github.com/rohitsinghlab/deltanmf.

## Acknowledgements

R.S. acknowledges the support of the Chan-Zuckerberg Initiative and Duke University’s Whitehead Scholarship. Some figures were made with biorender.com.

## A Supplementary Methods

### A.1 Stability and consensus initialization

Like standard NMF solvers, our GPU-based formulation also finds local optima. We found that the initialization of **W** and **H** parameters influenced the final **W**_**eff**_ and **H**_**eff**_ . Following cNMF, we do multiple short runs and our final reported parameters are from a DeltaNMF run initialized with consensus parameters **W**_**0**_ and **H**_**0**_:

#### 1. Multiple runs

Execute *R* independent factorizations (default *R* = 20) that each yield 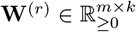. Each replicate uses our minibatch NMF with multiplicative updates (MU) for **H** and columwise MU for **W**. Replicates are deliberately short (default 100 epochs, much fewer than the final run) to sample diverse attraction basins rather than fully converge every run. We seed each replicate independently.

#### 2. Pooling and normalization

Collect all *R* × *k* program columns {**w**_*i*_} and *L*_2_-normalize each **w**_*i*_ ← **w**_*i*_*/*∥**w**_*i*_∥_2_. This puts programs on a common scale.

#### 3. Density filtering

For each **w**_*i*_, compute the mean distance to its *L* nearest neighbors (default *L* = ⌊0.3*R*⌋ ). Remove outliers whose local density exceeds the *q*-quantile (default *q* = 0.95). This quantile-based density filter adapts to datasets with different stability profiles and reduces spurious programs compared to fixed thresholds.

#### 4. Clustering and aggregation

Cluster the retained {**w**_*i*_} into *k* groups by *k*-means in the normalized space. Stack the cluster means to obtain **W**_0_, then refit **H**_0_ by multiplicative update against fixed **W**_0_ in minibatches. The final DeltaNMF run is initialized with (**W**_0_, **H**_0_) and trained for many more epochs with the full objective and regularizers.

### A.2 Stage 2 residual-based initialization

We initialize **W**_2_ for Stage 2 using a residual approach:

1. Project cases onto the fixed baseline to get estimated initial usages 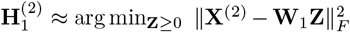. Initial usages usages are calculated from first computing the pseudoinverse and then fixed factor NMF computed via coordinate descent with **W**_**1**_ fixed. The resulting matrix captures the variance in **X**^(2)^ that can be explained by **W**_1_ learned on the control cells **X**^(1)^.
2. Form a nonnegative residual 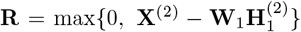. This corresponds to the remaining unexplained variance in **X**^(2)^ that cannot be explained by **W**_1_ learned on the control cells **X**^(1)^.
3. Run the consensus MU-based initialization on **R** to initialize 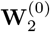 and 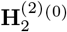.
4. Balance column norms by a scalar *s* so that the median *ℓ*_2_ norm of 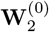 matches that of **W**_1_; set 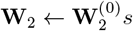.
5. Solve for initial joint usages **H**^(2)^ by coordinate descent on the combined program matrix [**W**_1_ **W**_2_].
6. Then perform Stage 2 neural optimization to refine **W**_2_ and **H**^(2)^ while keeping **W**_1_ fixed.

### A.3 Training details

We trained using the Adam optimizer, with softplus activation (*β* = 5.0) to enforce non-negativity. For all results, we used a learning rate of 0.01 and a maximum of 10,000 epochs. For NTC discovery, we first ran consensus NMF with 20 replicates (100 epochs each, density threshold quantile 0.95) to initialize **W** and **H** and then refined with foundation model regularization. To preserve the non-negativity required for NMF while normalizing gene variance, we applied a non-centering unit-variance scaling. For each gene *g*, we computed the standard deviation *σ*_*g*_ across all cells. The expression entries were scaled as *X*_*ig*_ ← *X*_*ig*_*/σ*_*g*_. This prevents high-expression housekeeping genes from dominating the Frobenius reconstruction error, ensuring that the optimization gives equal weight to variance explained in lower-expression signaling factors.

### A.4 Dataset Details

#### Coronary Artery Disease (CAD) Perturb-seq

We utilized the large-scale CRISPR interference (CRISPRi) screen described by Schnitzler et al. [19], performed in telomerase-immortalized human aortic endothelial cells (TeloHAEC). This cell line was selected for its relevance to arterial biology and validated by comparison to primary human right coronary artery endothelial cells. The library targeted 2,285 genes, specifically focusing on 1,661 expressed genes located within ±500kb of 228 *non-lipid* CAD GWAS signals (loci not associated with circulating lipids), to isolate mechanisms intrinsic to the vessel wall. Cells were harvested after 5 days of doxycycline induction to capture downstream transcriptional consequences. The final quality-controlled dataset comprises 214,449 singlet cells sequenced to a median depth of ∼10,870 UMIs per cell. For Stage 1 (baseline discovery), we restricted the analysis to the 5,506 cells that received non-targeting control (NTC) guides.

#### Alzheimer’s Disease (AD) Perturb-seq

We analyzed the CRISPR interference (CRISPRi) component of the screen described by Wang et al. [22]. This screen was performed in primary human macrophages (hM*Φ*) derived from peripheral blood mononuclear cells (PBMCs) to better model microglia physiology. The library targeted 203 genes, including 71 AD risk genes prioritized by GWAS and 132 genes linked to microglial function. We utilized the quality-controlled dataset of 28,446 cells, using the 2,184 cells receiving non-targeting guides to define the baseline macrophage manifold in Stage 1.

## A.5 Gene selection strategy

For comparisons to cNMF on simulated data in the Fig. 2 benchmarking, we match the preprocessing in Kotliar et al. by restricting to the 2,000 most highly variable genes (HVGs) based on overdispersion. For all downstream biological analyses (Figures 3–5), we retain all genes rather than restricting to HVGs. This follows recommendations from Schnitzler et al. that rare or perturbation-specific programs may depend on genes that fall outside the HVG set [19].

### A.6 Paired-Program Simulation Framework

To rigorously test the ability of DeltaNMF to disentangle simultaneously active gene programs, we extended the `scsim` framework [25] to support a “paired-program” architecture.

#### Generative Model

We simulated single-cell RNA-seq counts for *N* = 15, 000 cells and *G* = 25, 000 genes across *C* = 10 distinct cell groups. Each group *g* is associated with a unique pair of gene programs: a Control program (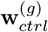) and a Case program (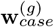).

Program gene sets were chosen from the background gene pool. For the specific simulation benchmark used in Figure 4:

- **Control Programs:** 600 genes each, enriched by fold-change *FC* = 3.0.
- **Case Programs:** 200 genes each, enriched by fold-change *FC* = 3.0.
- **Overlap:** The gene sets for paired Control and Case programs were disjoint (overlap fraction = 0.0), ensuring that any observed correlation between inferred factors is due to model misspecification (blending) rather than shared ground-truth genes.

#### Cellular States

Within each group, 50% of cells were assigned a “Control” label and 50% a “Case” label.

- **Control Cells** express only the group-specific control program: 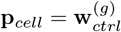
- **Case Cells** express a convex combination of both programs: 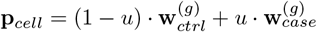

where *u* represents the disease program usage. We fixed *u* = 0.5 for all case cells to simulate a strong coupling. The final cell gene means were derived by mixing this program profile **p**_*cell*_ with a group-specific background expression profile (mixing coefficient *β* = 0.7).

#### Technical Noise

We applied realistic technical noise models consistent with the Splatter framework used by Kotliar et al. [12], including:

- Library size variation (log-normal, *µ* = 7.64, *σ* = 0.78).
- Mean-variance trend (BCV dispersion = 0.448).
- Doublet simulation (1,000 doublets added).
- Dropout and outlier gene expression.

Raw counts were sampled from a Poisson distribution parameterized by these adjusted means.

### A.7 Identification of program regulators

To identify the upstream genetic drivers (regulators) of inferred gene programs, we leveraged the Perturb-seq experimental design, which provides a mapping between cells and the specific CRISPR guides they received. Let **H** ∈ ℝ^*k*×*n*^ be the inferred usage matrix (where *k* is the number of programs and *n* is the number of cells) and **G** ∈ {0, 1} ^*n*×*g*^ be the binary guide assignment matrix (where *g* is the number of unique guide sequences).

First, we computed the average activity of each program *p* in cells containing guide *j*:

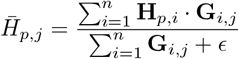

where the denominator represents the number of cells receiving guide *j* (with *ϵ* added for numerical stability).

Since multiple distinct guide sequences may target the same gene, we aggregated these guide-level scores to the gene level. For each target gene *t*, we identified the set of guides 𝒥_*t*_ targeting that gene and computed the mean program activity:

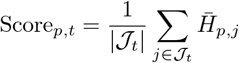

We then ranked target genes by this score for each program. The top 15 target genes with the highest mean program activity were defined as the candidate regulators for that program.

## B Supplementary Results

**Fig. B.1:**
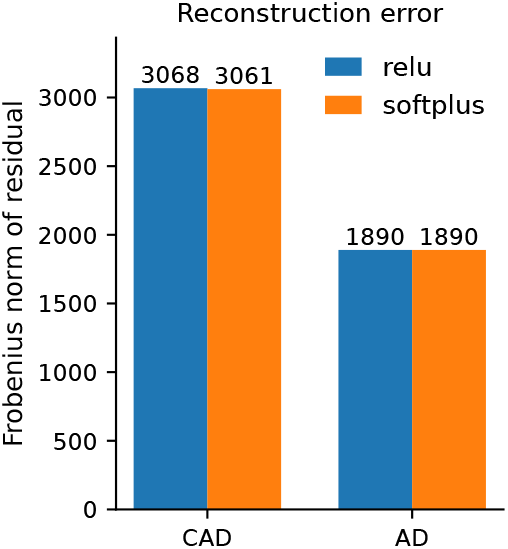
Softplus activation improves convergence stability compared to ReLU.

**Fig. B.2:**
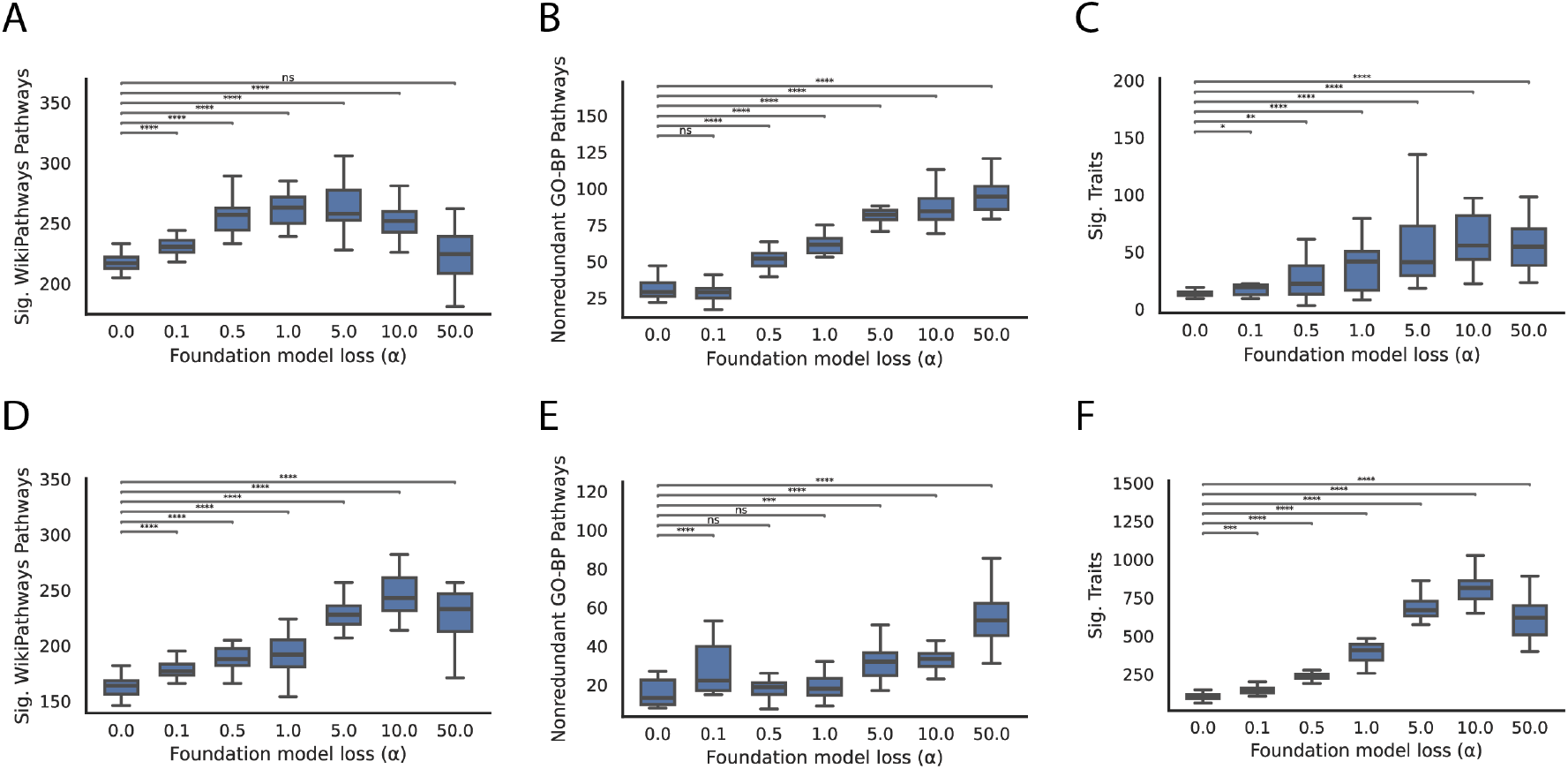
Extended biological coherence metrics for scGPT-regularized DeltaNMF. Additional quantification of program quality for the CAD (top row) and AD (bottom row) Perturb-seq datasets using scGPT regularization (*α* sweep from 0 to 50, applied during final 200 epochs). **(A, D)** Enrichment for WikiPathways, showing the number of significant pathways (*FDR* < 0.05) associated with learned programs. **(B, E)** Number of non-redundant GO Biological Process terms per program after collapsing semantically similar terms, indicating sharper functional delineation. **(C, F)** Trait relevance measured by enrichment for GWAS-implicated genes (Open Targets L2G), showing that regularization recovers programs with stronger links to known traits and disease biology. Data are presented as mean ± s.d. across 20 bootstrap replicates.

**Fig. B.3:**
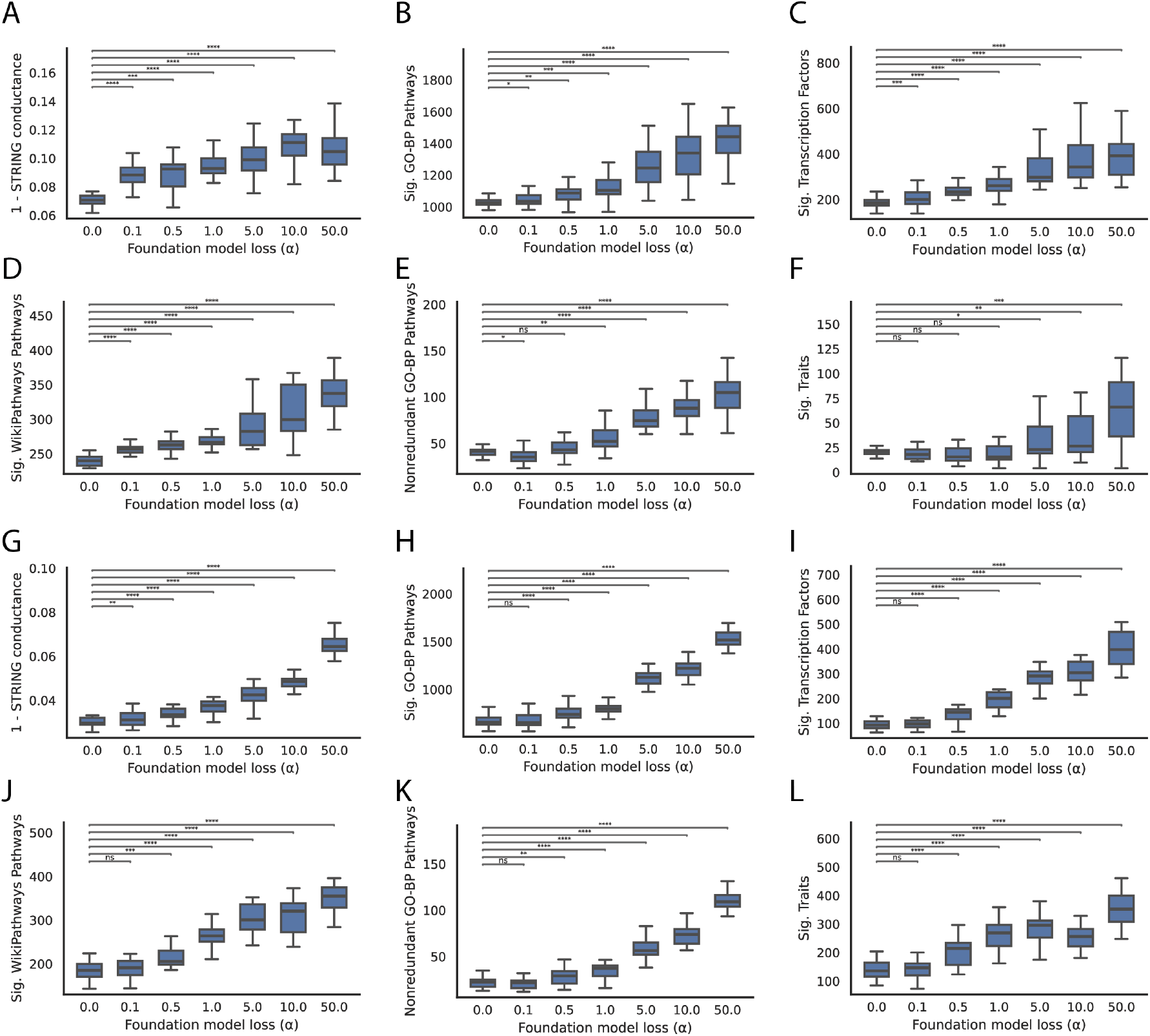
Foundation model regularization improves program coherence using TranscriptFormer embeddings. Validation of the DeltaNMF regularization strategy using an alternative foundation model. The analysis from Fig. 3 was repeated using gene-gene similarity graphs constructed from TranscriptFormer embeddings rather than scGPT. **(A-F)** Evaluation on the CAD Perturb-seq dataset showing improvements in STRING network cohesion, GO-BP enrichment, and TRRUST transcription factor enrichment as a function of regularization strength *α*. **(G-L)** Evaluation on the AD Perturb-seq dataset showing consistent trends. The similarity in performance improvements suggests that the regularization benefits derive from general transcriptomic co-activity patterns captured by large-scale transformers rather than model-specific artifacts.

**Fig. B.4:**
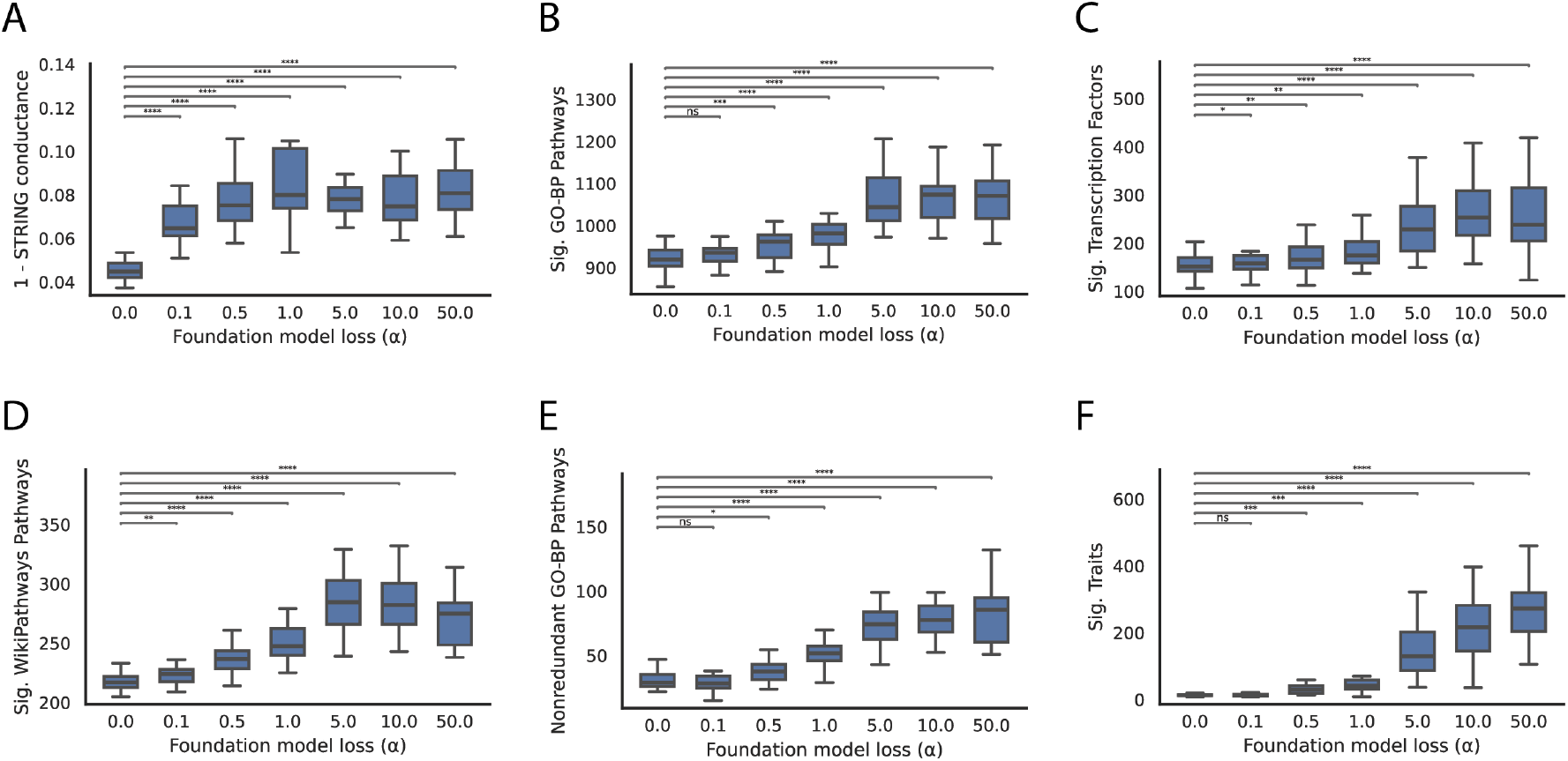
Effect of extended regularization schedule on CAD program coherence. Evaluation of program coherence when scGPT-based graph Laplacian regularization is applied for a longer duration. Here, regularization was applied for the final 1000 epochs of training (compared to 200 epochs in the main analysis) on the CAD Perturb-seq dataset.

**Fig. B.5:**
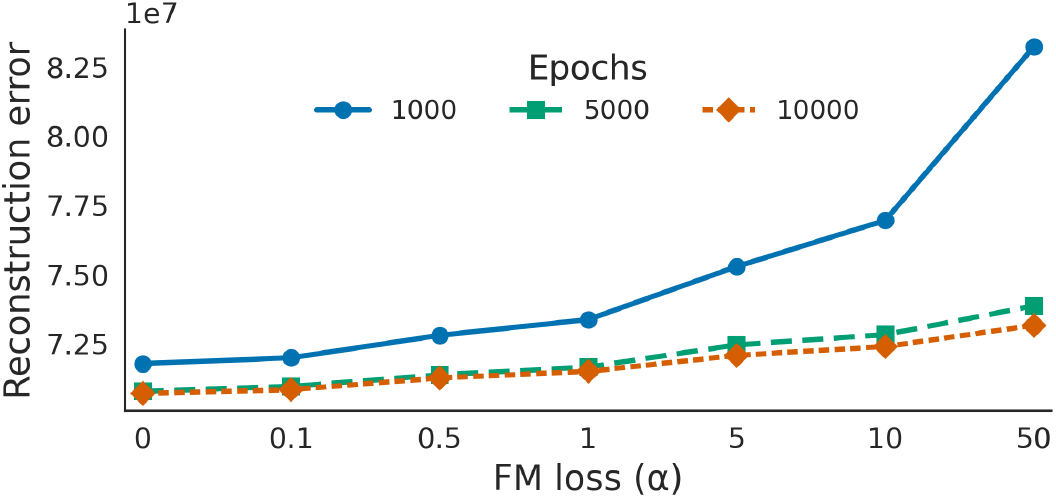
Stability of reconstruction error across regularization strengths. Assessment of the trade-off between biological coherence and data fidelity. The plot displays the final reconstruction error (normalized Frobenius loss) on the CAD Perturb-seq dataset across the full sweep of regularization strengths (*α* ∈ [0, 50]).

**Fig. B.6:**
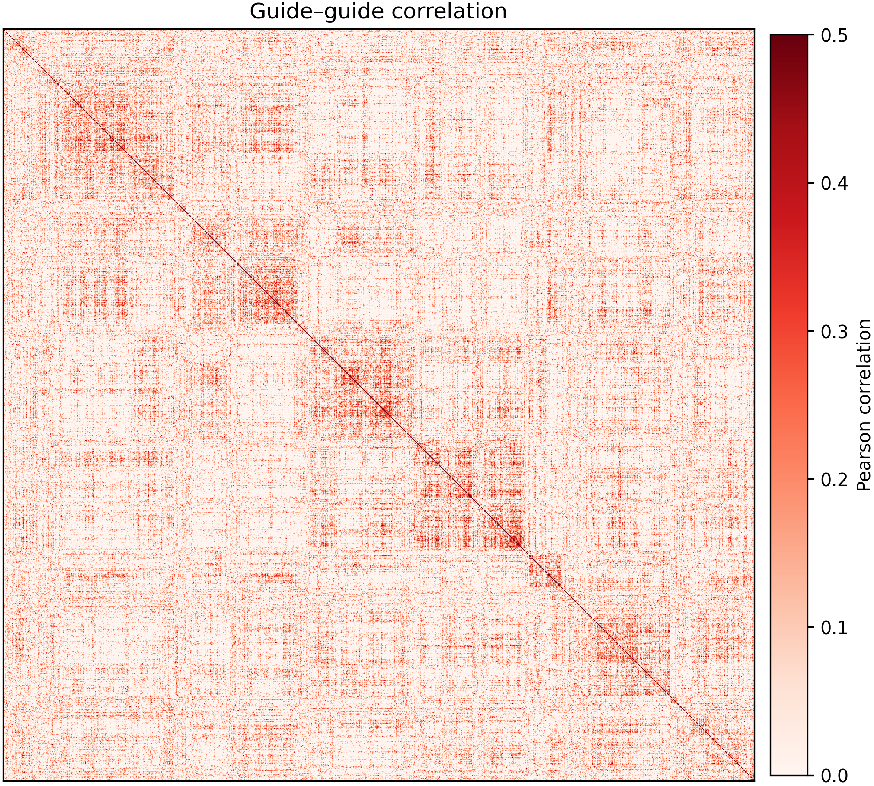
Guide–guide Pearson correlation matrix for CAD case cells, computed from DeltaNMF program usages and clustered by correlation distance. Blocks indicate groups of perturbations that converge on similar transcriptional programs.

**Fig. B.7:**
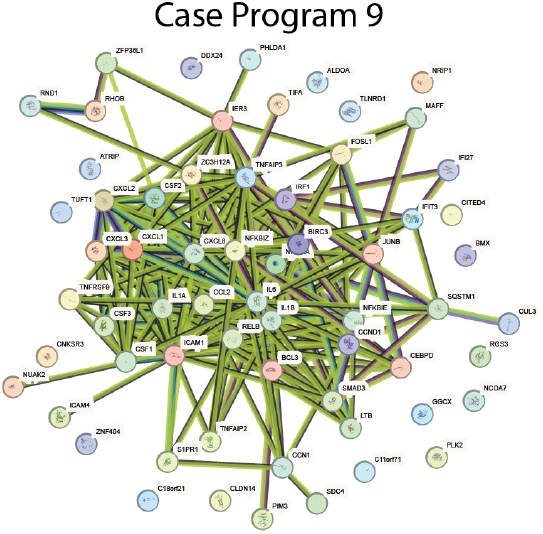
STRING pathway prioritized by MAGMA for Case Program 9.

**Fig. B.8:**
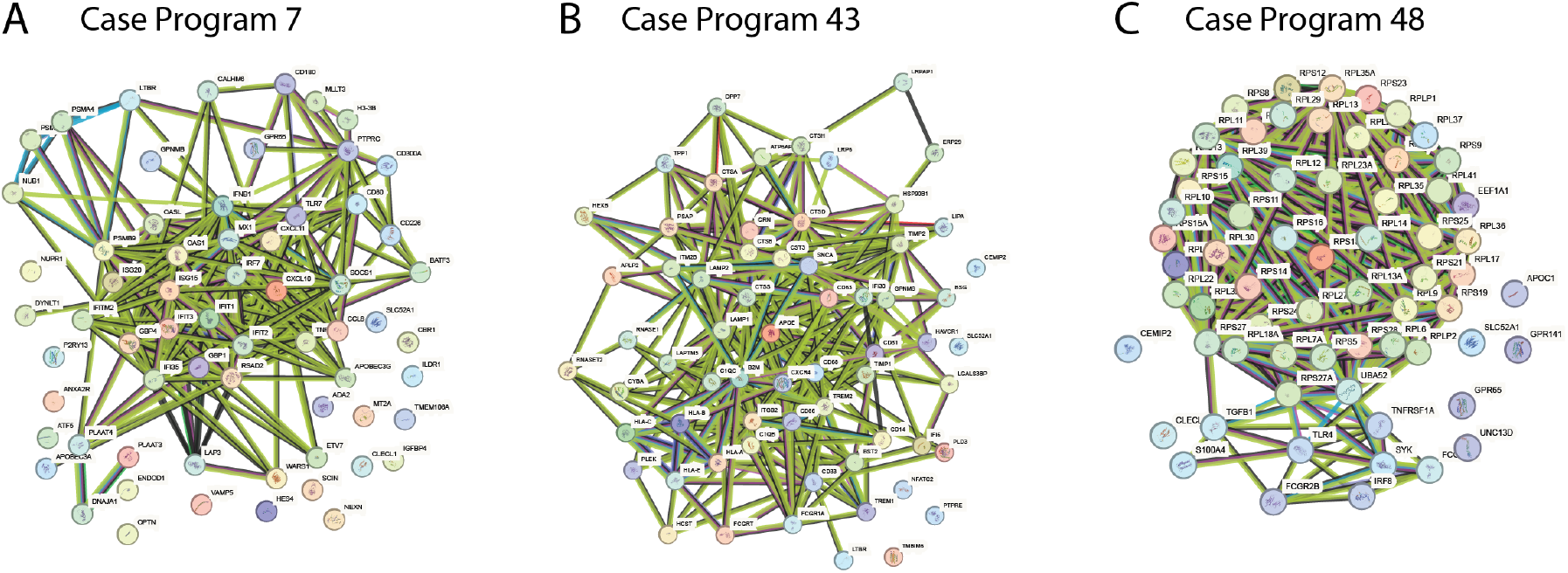
DeltaNMF recovers distinct, biologically coherent macrophage states associated with Alzheimer’s disease risk as visualized by STRING network visualizations of the top 50 genes and top 15 regulators from three key case programs (programs 7 (A), 43 (B), and 48 (C)).

